# Taste receptors’ profiling in glioblastoma

**DOI:** 10.1101/2025.06.24.661264

**Authors:** Ana R. Costa, Ana C. Duarte, Isabel Gonçalves, Catarina L. Ferreira, J. Eduardo Cavaco, Isidro Ferrer, Tomomi Furihata, Helena Marcelino, José F. Cascalheira, Cecília R.A. Santos

## Abstract

Glioblastoma is the most common and aggressive form of primary brain cancer. Despite significant progress in the development of promising therapeutic agents, cancer-targeting therapies often fail to achieve effective concentrations in the brain, limiting their therapeutic efficacy. As such, a deeper understanding of how glioblastoma tumours interact with their microenvironment and assess the chemical composition therein can reveal novel therapeutic strategies. Recent studies have highlighted the critical role of taste receptors, particularly bitter taste receptors (TAS2Rs) and their ligands in cancer progression and metastasis. Activation of TAS2Rs by both natural and synthetic compounds has been associated with drug resistance, apoptosis, and the proliferation of malignant tumours. The sweet taste receptor TAS1R2/TAS1R3, or the umami receptor TAS1R1/TAS1R3, as recognised sensors of glucose levels and aminoacids, respectively, are also of interest in the context of cancer metabolism. In this study, we investigated the expression and function of TAS2Rs, TAS1R2/TAS1R3 and TAS1R1/TAS1R3 and all the taste signalling pathway machinery in glioblastoma. Our findings demonstrate that TAS1R2/TAS1R3, TAS1R1/TAS1R3, and 20 out of the 26 human TAS2Rs are present and active in glioblastoma cells, with expression differences in glioblastoma cells and human tumours. The expression of such a large number of members of this family of receptors, highlight the significance of the taste transduction pathway in this form of brain cancer, where these receptors might be essential for the crosstalk between glioblastoma and its microenvironment.

## 1. Introduction

Glioblastoma is the most common and aggressive form of primary brain cancer, classified grade 4 by the World Health Organization, with a 5-year survival of approximately 3% and a median survival of less than 18 months [1,2]. Despite the impressive advances in the development of promising drugs for the treatment and/or diagnosis of tumours in the central nervous system (CNS), glioblastomas remain resistant to pharmacologic interventions. This is a consequence of chemo- and radioresistant cancer stem-like cells relapsing, adaptation to changes in the tumour microenvironment [3], and the highly infiltrative nature of the tumours that impair their complete resection [4–6].

The human taste receptor family is composed of 26 bitter taste receptors (TAS2R), the sweet taste receptor (TAS1R2/TAS1R3), the umami receptor (TAS1R1/TAS1R3), and the ionotropic sour and the salt receptors. Ionic taste stimuli are detected through ion channels, with the proton-gated channel Otop-1 acting as the primary receptor for sour taste, while the epithelial sodium channel (ENaC) functions as the receptor for salty taste. In contrast, the receptors responsible for sweet, umami, and bitter taste belong to the superfamily of G protein-coupled receptors (GPCRs). The three TAS1R genes encode heterodimers that form the key umami receptor (TAS1R1/TAS1R3) and the sweet taste receptor (TAS1R2/TAS1R3), whereas the 26 functional bitter taste receptors are part of the TAS2R gene family [8]. These mediate signal transduction in response to a wide range of specific bitter compounds [9,10]. In the canonical pathway, taste receptor activation leads to the dissociation of heterotrimeric G protein into α-gustducin and Gβ3/Gγ13 subunits, which in turn activate downstream phospholipase-C β2 that hydrolyses phosphatidylinositol-4,5-bisphosphate to inositol-1,4,5-triphosphate (IP3) and diacylglycerol. IP3 triggers an increase in intracellular Ca^2+^ levels that, in turn, activate the transient receptor potential cation channel subfamily M member 5, causing cell depolarization [11,12]. Apart from sensing compounds in the oral cavity, taste receptors were also described in several extraoral tissues, like the gastrointestinal tract, airway epithelium, heart, testis and brain [13–16], and have also been implicated in a diversity of metabolic and developmental pathologies such as diabetes, schizophrenia, Parkinson’s disease and various types of cancer [17–20]. Experimental and clinical data revealed that taste receptors and their ligands have a role in cancer progression and metastasis by controlling many aspects of tumorigenesis, proliferation, migration, cancer cell invasion and cancer-related signalling pathways [21,22]. Also, expression differences between cancer and normal cells are frequently found [23]. However, taste receptor expression and possible chemosensory function(s) in glioblastoma cells have not been studied. Given the potential of these receptors in the crosstalk of glioblastoma cells with their microenvironment, we first analysed the expression of sweet, *umami*, and bitter receptors and compared their expression between a human astrocyte and three different glioblastoma cell lines. Bitter taste receptors TAS2R4, TAS2R5, TAS2R14 and TAS2R39, and the sweet taste receptor TAS1R2/TAS1R3, were also analysed in human glioblastoma samples.

## 2. Materials and methods

### 2.1 Materials

Probenecid (#P8761; CAS No 57-66-9), a known bitter taste receptors’ antagonist, and denatonium benzoate (#D5765; CAS No 3734-33-6) were obtained from Sigma-Aldrich. Probenecid was dissolved in 1 N NaOH at 0.17 M and diluted in Tyrode’s solution or culture medium. Denatonium benzoate was prepared in dimethyl sulfoxide (DMSO) and freshly dissolved in Tyrode’s solution or culture medium before the experiments. FURA-2AM (#F1221) and pluronic acid F-127 were purchased from ThermoFisher Scientific.

The RT-PCR and RT-qPCR primers (Supplementary Tables S1 and S2) were obtained from STAB VIDA, and the primary and secondary antibodies (Supplementary Table S3), previously validated, were purchased from Santa Cruz Biotechnology, ThermoFisher Scientific and Sigma-Aldrich.

### 2.2 Cell Culture

The conditionally immortalised human astrocyte cell line HASTR/ci35 [24] was developed by one of the co-authors (T. Furihata) at Chiba University (Japan). HASTR/ci35 cells were cultured at 33 °C, 5% CO_2_ using the recommended Astrocyte Medium (Gibco™ #A1261301) supplemented with 1% N2 supplement, 10% (v/v) fetal bovine serum (FBS), penicillin (100 IU/mL)/streptomycin (100 μg/mL) and 4 μg/mL blasticidin S (Abcam #ab141452).

Human malignant glioblastoma cell lines U-87MG and U-373MG were kindly provided by Dr. Joseph Costello (University of California, San Francisco), and SNB-19 was obtained from the German Collection of Microorganisms and Cell Cultures. Glioblastoma cells were grown in Dulbecco’s modified Eagle’s medium (DMEM) high glucose with stable glutamine (bioWest #L0103) supplemented with 10% (v/v) FBS and penicillin (100 IU/mL)/streptomycin (100 μg/mL), and incubated in a humidified atmosphere containing 5% CO_2_ at 37 °C.

### 2.3 Detection of taste receptor transcripts in human astrocytes and glioblastoma cell lines by RT-PCR

Total RNA was isolated from glioblastoma cells using TRI Reagent^®^ (Sigma Aldrich #T9424), following the manufacturer’s instructions. The RNA concentration was determined by OD measurement at 260 nm, and the quality of RNA was evaluated by agarose gel electrophoresis. cDNA was synthesised using NZY M-MuLV Reverse Transcriptase (NZYTech #MB083) according to the protocol supplied by the manufacturer. For the RT-PCR, cDNA was amplified by HOT FIREPol^®^ DNA Polymerase (Solis BioDyne #01-02) and specific primers (Supplementary Table S1) in a final volume of 10 µL. The RT-PCR protocols comprised an initial denaturation for 3 minutes at 95°C, followed by 40 cycles of denaturation at 94°C (30 seconds), annealing (30 seconds) and elongation at 72°C (15-30 seconds), and 5 minutes of final elongation at 72°C. The PCR products were analysed by electrophoresis on 1% or 1.5% agarose gels and visualised using GreenSafe Premium (NZYTech #MB13201) staining. In addition, to confirm the sequence identity, RT-PCR products were purified, and Sanger sequenced by STAB VIDA (Portugal).

For real time PCR and protein expression analysis, we chose bitter receptors TAS2R4, TAS2R5, TAS2R14 and TAS2R39, and sweet taste receptor TAS1R2/TAS1R3 based on the following criteria: 1) validated expression by RT-PCR; 2) number and therapeutic relevance of known ligands; 3) primary antibodies commercially available, suitable for both Western blot and immunofluorescence techniques; 4) previous validation of primary antibodies. In addition, TAS2R4 and TAS2R14 have previously been associated with regulation of cancer cell migration and proliferation [25], while TAS2R5 and TAS2R39 have well-characterized ligands and validated antibodies for reliable detection.

### 2.4 Analysis of the mRNA expression levels of TAS2R by real-time quantitative RT-PCR

Analysis of the mRNA expression levels of TAS2R4, TAS2R5, TAS2R14, TAS2R39 and the housekeeping gene glyceraldehyde 3-phosphate dehydrogenase (GAPDH) were performed by RT-qPCR using the NZYSpeedy qPCR Green Master Mix (NZYTech #MB224). A validation assay was previously performed with cDNA serial dilutions for all the genes. Cycling conditions were the following: 95 °C for 2 minutes, followed by 40 cycles of 95 °C for 5 seconds, 58 °C for 10 seconds, and 72 °C for 3 seconds. Primer sequences for TAS2Rs and GAPDH used for RT-qPCR are shown in Supplementary Table S2. The ΔCt was calculated using GAPDH mRNA as the reference gene, and the ΔΔCt was calculated between the normalised ΔCt values from each glioblastoma cell line and the average Ct value from the human astrocytes cell line. The mRNAs’ relative expressions were calculated by the 2^-ΔΔCt^ method [26]. A melting curve analysis was performed after the final cycle to ensure a single product was obtained.

### 2.5 Detection of taste receptors and GNAT3 protein in human astrocytes and glioblastoma cell lines

#### 2.5.1 Western blot analysis

Total protein extracts were prepared by suspending glioblastoma cells in phosphate-buffered saline (PBS 1x), followed by centrifugation at 10 000 g for 7 min at 4 °C, and homogenisation in RIPA buffer [150 mM NaCl, 0.5% Sodium deoxycholate, 0.1% SDS, 1% Triton X-100, 50 mM Tris pH 8.0, 1 mM PMSF and 10 µL/mL of complete EDTA Free protease inhibitor cocktail (Roche #11873580001)]. Total protein measurement was performed with Pierce BCA Protein Assay Kit (ThermoFisher Scientific #23227) following the manufacturer’s instructions. An amount of 50 µg of total protein was mixed with a loading buffer containing 4% β-mercaptoethanol, followed by denaturation for 5 min at 100 °C, and then loaded in a 12.5% SDS-PAGE. Total proteins were transferred to a PVDF membrane 0.45 μm (GE Healthcare #10600023), which was blocked with 5% non-fat dry milk in Tris-buffered saline (TBS 1x), for 1h at room temperature (RT). After overnight incubation at 4 °C with the primary antibodies GNAT3, bitter taste receptors TAS2R4, TAS2R5, TAS2R14 and TAS3R39, and sweet taste receptor TAS1R2/TAS1R3 (Supplementary Table S3), membranes were rinsed three times with TBS containing 0.1% of Tween-20 (TBS-T) and incubated with the respective HRP-linked secondary antibody for 1 h at RT. Moreover, GNAT3 primary antibody specificity was assessed through parallel incubation with the respective peptide. After this, signal detection was performed with SuperSignal™ West Pico PLUS Chemiluminescent Substrate (ThermoFisher Scientific #34577), and images were acquired with the Image Lab software in a ChemiDoc™ MP (Bio-Rad). This experiment was done with at least three different cell passages. Additionally, expression of GNAT3 and taste receptors was normalised with β-actin incubated for 1 h at RT before incubation for 1 h with HRP-conjugated goat anti-mouse secondary antibody. Protein bands were quantified using the Image Lab software (Bio-Rad).

#### 2.5.2 Immunocytochemistry

The presence of GNAT3, bitter taste receptors TAS2R4, TAS2R5, TAS2R14 and TAS2R39, and sweet taste receptor TAS1R2/TAS1R3 was investigated in glioblastoma cell lines by fluorescence immunocytochemistry. Briefly, cells were seeded and grown on glass coverslips until 60-70% confluence. After removing the medium, cells were fixed with paraformaldehyde (PFA) 4% for 10 min followed by 1 h blocking in PBS containing 3% bovine serum albumin (BSA) and 0.2% Triton X-100 at RT. Cells were incubated overnight at 4 °C with the primary antibodies (Supplementary Table S3) in a blocking solution. In negative controls, the primary antibody was omitted. Glioblastoma cells were then incubated for 1h with Alexa Fluor^®^ 488 goat anti-rabbit IgG conjugate or Alexa Fluor^®^ 647 chicken anti-goat IgG conjugate. Finally, coverslips were incubated for 10 min with the fluorescent dye Hoechst 33342 to visualize their nuclei. After several washes, cells were mounted onto microscope slides and visualized under a confocal microscope LSM 710 (Zeiss) using a magnification of 63x (Plan-Apochromat 63x/1.4 Oil DIC M27).

### 2.6 Detection of taste receptors in human samples of glioblastoma by immunohistochemistry

Cases for the immunohistochemistry study were obtained from the Institute of Neuropathology Brain Bank (HUB-ICO-IDIBELL Biobank) following the guidelines of the Spanish legislation on this matter (Real Decreto 1716/2011) and the approval of the local ethics committee of the Bellvitge University Hospital-IDIBELL. Human glioblastoma samples were fixed in buffered formalin for no less than 3 weeks and then embedded in paraffin. Paraffin-embedded human glioblastoma slices, from men (n=4) and women (n=6), were pre-treated with Trilogy™ (Cell Marque™ #920P) which combines deparaffinisation, rehydration and unmasking, following manufacturer’s recommendations. After washing with TBS-T, endogenous peroxidases activity was blocked with 3% H_2_O_2_ for 10 min at RT. Slices were then washed twice with TBS-T. Next, slices were incubated for 1 h at RT with the following primary antibodies TAS2R4, TAS2R5, TAS2R14, TAS2R39, TAS1R2 and TAS1R3 (Supplementary Table S3). Slices were washed twice with TBS-T and treated with HiDef Detection™ HRP Polymer System (Cell Marque™ #954D). First, HiDef Detection™ Amplifier was applied in the human glioblastoma slices for 10 min RT, washed twice with TBS-T, followed by HiDef Detection™ HRP Polymer Detector also for 10 min at RT. After slices washing with TBS-T, immunoreactivity was detected with diaminobenzidine (DAB) for 10 min RT. Slices were washed twice with TBS-T, and tissue sections were stained with Mayer’s haematoxylin for 3 min RT to allow nuclei visualisation. Negative control slices were treated under the same conditions without primary antibody. After dehydration, the slices were mounted with Q Path^®^ Coverquick 2000 (VWR #805547530) and the images were acquired in a Widefield Axio Imager Z2 (Zeiss) using a 20x magnification (Plan-Apochromat 20x/0.8 M27).

### 2.7 Single-cell calcium imaging functional assay

Functional analysis was assessed by analysing Ca^2+^ cell responses to the well-known bitter compound denatonium benzoate (DnB). This single-cell Ca^2+^ imaging assay was performed in the presence and absence of the TAS2Rs antagonist probenecid in glioblastoma U-373MG cells. In brief, approximately 3.5x10^4^ glioblastoma cells were seeded in μ-slide 8 well ibiTreat (Ibidi #80826) and grown until 60-70% confluency, followed by measurement of changes in intracellular Ca^2+^ levels after stimulation. Glioblastoma cells were loaded with 5 μM of FURA-2 AM and 0.02% pluronic acid F-127 in culture medium for 45 min. Next, cells were washed twice with Tyrode’s solution pH 7.4 [NaCl 140 mM, KCl 5 mM, MgCl_2_ 1 mM, CaCl_2_ 2.0 mM, Na-pyruvate 10 mM, Glucose 10 mM, HEPES 10 mM, NaHCO_3_ 5 mM] and loaded with Tyrode’s solution for 30 min. After that, dose-response experiments were performed with a range of DnB concentrations (2.5 and 5 mM), in the presence or absence of 30 min incubation with probenecid (1 mM). The μ-slide plates were placed on a Widefield Axio Observer Z1 inverted microscope (Zeiss). Stock solution of DnB and probenecid was freshly prepared in Tyrode’s solution before the experiments. The stimulus was applied manually with a micropipette after the baseline was recorded. The intracellular Ca^2+^ levels were evaluated by quantifying the ratio of the fluorescence emitted at 520 nm following alternate excitation at 340 nm and 380 nm, using a Lambda DG4 apparatus (Sutter Instrument) and a 520 nm bandpass filter (Zeiss) under a Fluar 40x/1.30 Oil M27 objective (Zeiss). Data were processed using the Fiji software (MediaWiki). Changes in fluorescence ratio (F=F340/F380) were measured in at least 20 cells, in three or more independent experiments. The response intensity, or intracellular Ca^2+^ variations ΔF/F0=(F-F0)/F0, where F0 corresponds to fluorescence ratio average at baseline (2 min acquisition before stimulus) and F correspond to maximum peak of fluorescence ratio evoked by the stimulus applied to the cells, was calculated.

### 2.8 Data Analysis

Statistical analysis and dataset comparisons were performed using GraphPad Prism 9.3.1 (GraphPad Software). Statistical significance was determined by One-Way ANOVA followed by the software’s recommended multiple comparisons post-hoc test. Results are presented as mean ± SEM of at least three independent experiments, and data were considered statistically different for a p-value<0.05.

## 3. Results

### 3.1 Human astrocytes and glioblastoma cells differentially express taste receptors

The bitter taste receptors (TAS2Rs) are conserved among mammals, with the human gene family comprising about 26 members [7,9]. The expression profile of TAS2Rs in a human astrocytes cell line and glioblastoma cells was assessed by RT-PCR using specific primers (Supplementary Table S1). The results demonstrated the mRNA expression of 20 TAS2Rs (1, 3, 4, 5, 7, 8, 9, 10, 14, 16, 38, 39, 40, 41, 42, 43, 44, 48, 50, and 60) (Table 1). No TAS2R13 mRNA was detected in any of the cells studied. The mRNA expression of TAS2R20/49, 30/47, 45 and 46 was not analysed because they share high levels of homology, impairing the design of specific primers. Interestingly, the observed patterns of TAS2Rs expression differed between the cell lines studied: for example, TAS2R60 was detected in all glioblastoma cells but not in human astrocytes (Table 1). Both specific subunits of the *umami* and sweet taste receptor, TAS1R1 and TAS1R2, respectively, as well as the shared subunit TAS1R3, were detected in all the cell lines (Table 1).

**Table 1.** mRNA expression profile of taste receptors in human astrocytes (HA) and glioblastoma (U-87MG, SNB-19, U-373MG) cell lines. RT-PCR was performed with cDNA synthesized from total RNA. The identities of the amplified products were confirmed by Sanger sequencing.

| Gene Symbol | HA | U-87MG | SNB-19 | U-373MG |
| --- | --- | --- | --- | --- |
| TAS1R1 | + | + | + | + |
| TAS1R2 | + | + | + | + |
| TAS1R3 | + | + | + | + |
| TAS2R1 | - | - | + | - |
| TAS2R3 | + | + | + | + |
| TAS2R4 | + | + | + | + |
| TAS2R5 | + | + | + | + |
| TAS2R7 | + | + | + | + |
| TAS2R8 | + | + | + | + |
| TAS2R9 | + | + | + | + |
| TAS2R10 | + | + | + | + |
| TAS2R13 | - | - | - | - |
| TAS2R14 | + | + | + | + |
| TAS2R16 | + | - | + | + |
| TAS2R38 | - | + | + | - |
| TAS2R39 | + | + | + | + |
| **TAS2R40** | + | + | + | + |
| **TAS2R41** | + | + | + | + |
| **TAS2R42** | + | + | + | + |
| **TAS2R43** | + | + | + | + |
| **TAS2R44** | + | + | + | + |
| **TAS2R48** | + | + | + | + |
| **TAS2R50** | + | - | + | - |
| **TAS2R60** | - | + | + | + |

To assess differences in the TAS2Rs mRNA expression between human astrocytes and glioblastoma cells, we performed RT-qPCR for TAS2R4, TAS2R5, TAS2R14, and TAS2R39. Compared to human astrocytes, TAS2R4 expression (Figure 1A) is significantly higher in SNB-19 and U-373MG cells (2.92- and 2.89-fold, respectively), and TAS2R5 expression (Figure 1B) is significantly higher in U-373MG cells (6.36-fold). TAS2R39 mRNA expression is significantly higher in U-87MG cells (Figure 1D) in comparison to human astrocytes (2.11-fold), SNB-19 (1.81-fold) and U-373MG (3.38-fold) glioblastoma cells. No significant differences were observed for TAS2R14 mRNA expression between astrocytes and cancer cell lines (Figure 1C).

**Figure 1.**
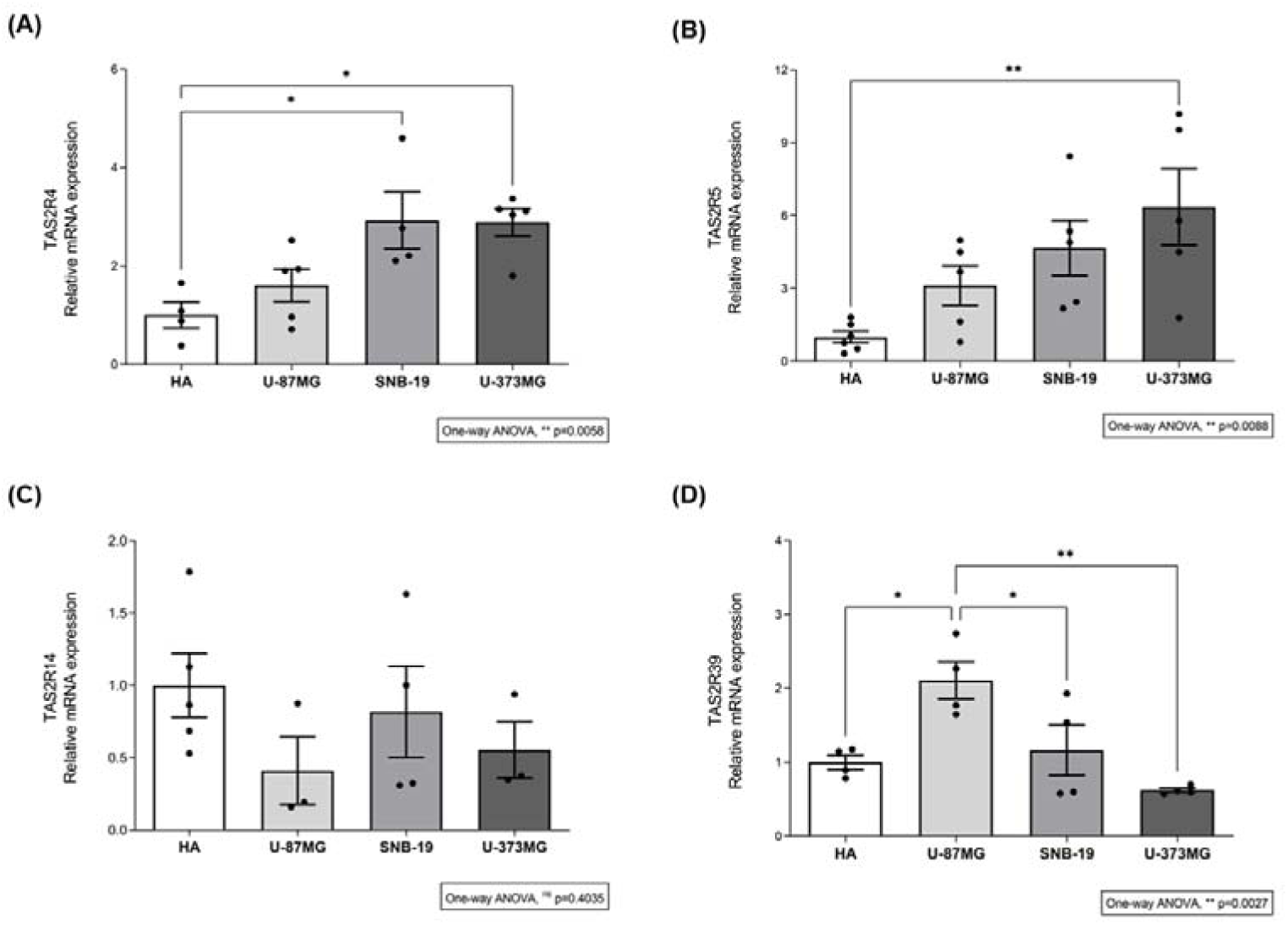
The mRNA of bitter taste receptors is differentially expressed in human astrocytes and in human glioblastoma cell lines. The mRNA expression levels of bitter taste receptors (A) TAS2R4, (B) TAS2R5, (C) TAS2R14 and (D) TAS2R39 normalized to the expression of the housekeeping gene GAPDH, were compared between human astrocytes (HA) and glioblastoma (U-87MG, SNB-19, U-373MG) cell lines by RT-qPCR. Results are presented as mean ± SEM and analysed by one-way ANOVA followed by Tukey’s multiple comparisons test. [N≥4 independent experiments; nsno significant; *p<0.05 and **p<0.01].

### 3.2 Human astrocytes and glioblastoma cells differentially express taste receptor proteins

We assessed the taste-related downstream effector protein α-gustducin (GNAT3), the four selected TAS2Rs, and the sweet taste receptor subunits TAS1R2/TAS1R3 in protein extracts of human astrocytes and glioblastoma cells by Western blot (Figure 2) and immunofluorescence (Figure 3) using available antibodies, previously validated by our research group [14]. The pre-incubation of the GNAT3 antibody with the respective peptide reduced the signal obtained, demonstrating the antibody specificity (Figure 2A).

**Figure 2.**
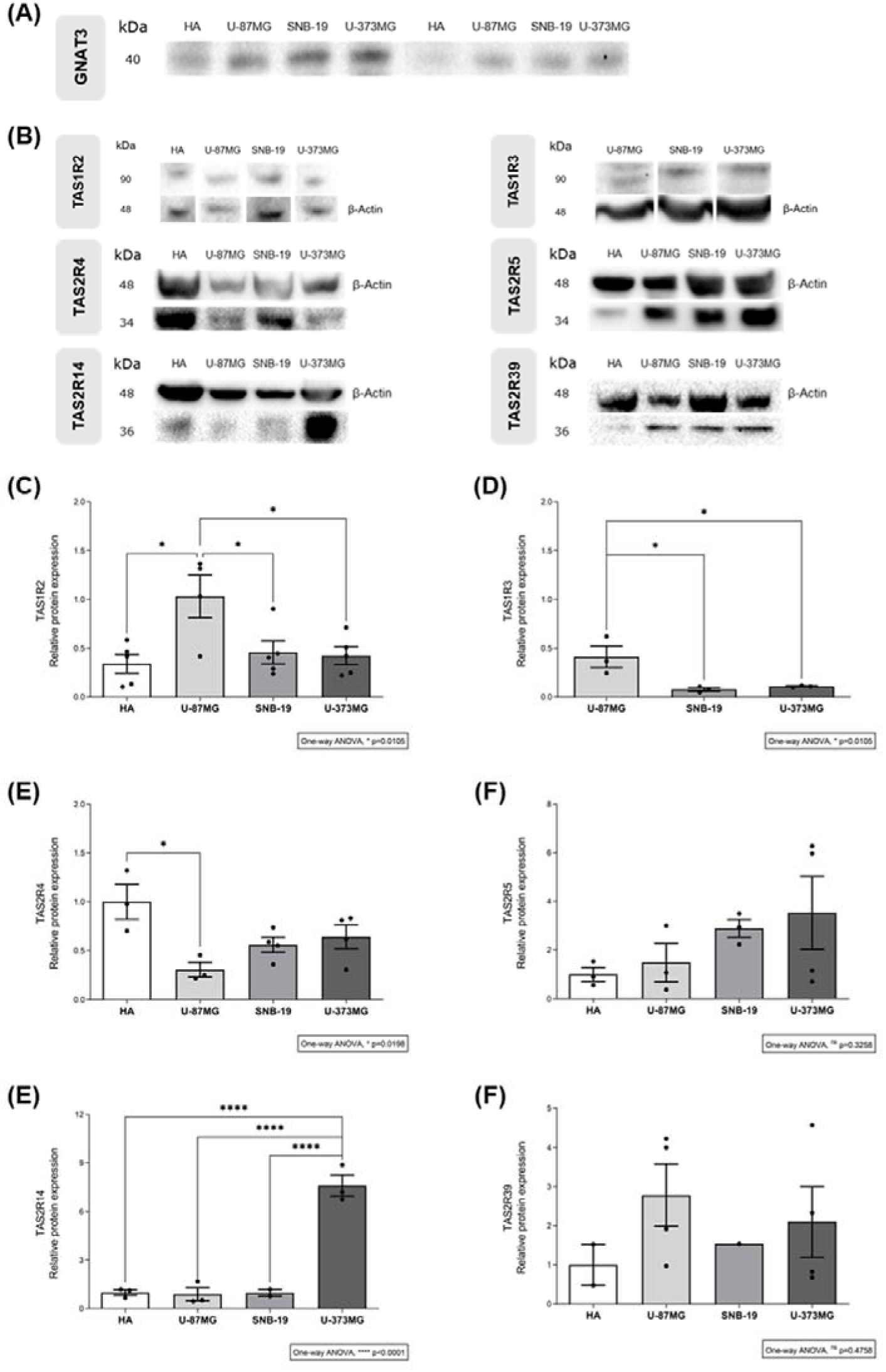
Sweet and bitter taste receptors, and taste signalling pathway effector GNAT3, are expressed in human astrocytes and glioblastoma cell lines. (A) GNAT3 expression was detected in all glioblastoma cell lines (left panel); when the GNAT3 antibody was pre-incubated with the respective peptide, the intensity of the immune reaction was strongly diminished showing the antibody specificity (right panel). (B) Western blot detection of sweet taste receptor subunits TAS1R2 and TAS1R3, and bitter taste receptors TAS2R4, TAS2R5, TAS2R14 and TAS2R39 in protein extracts of human astrocytes (HA) and glioblastoma (U-87MG, SNB-19, U-373MG) cell lines (representative images). (C–H) Relative quantification of the sweet taste receptor subunits TAS1R2 and TAS1R3, and bitter taste receptors TAS2R4, TAS2R5, TAS2R14 and TAS2R39 protein levels in HA and glioblastoma cells, normalized to β-actin levels. Results are presented as mean ± SEM and analysed by one-way ANOVA followed by Tukey’s multiple comparisons test. [N≥3 independent experiments; nsno significant; *p<0.05 and ****p<0.0001]. kDa – kilo Dalton.

**Figure 3.**
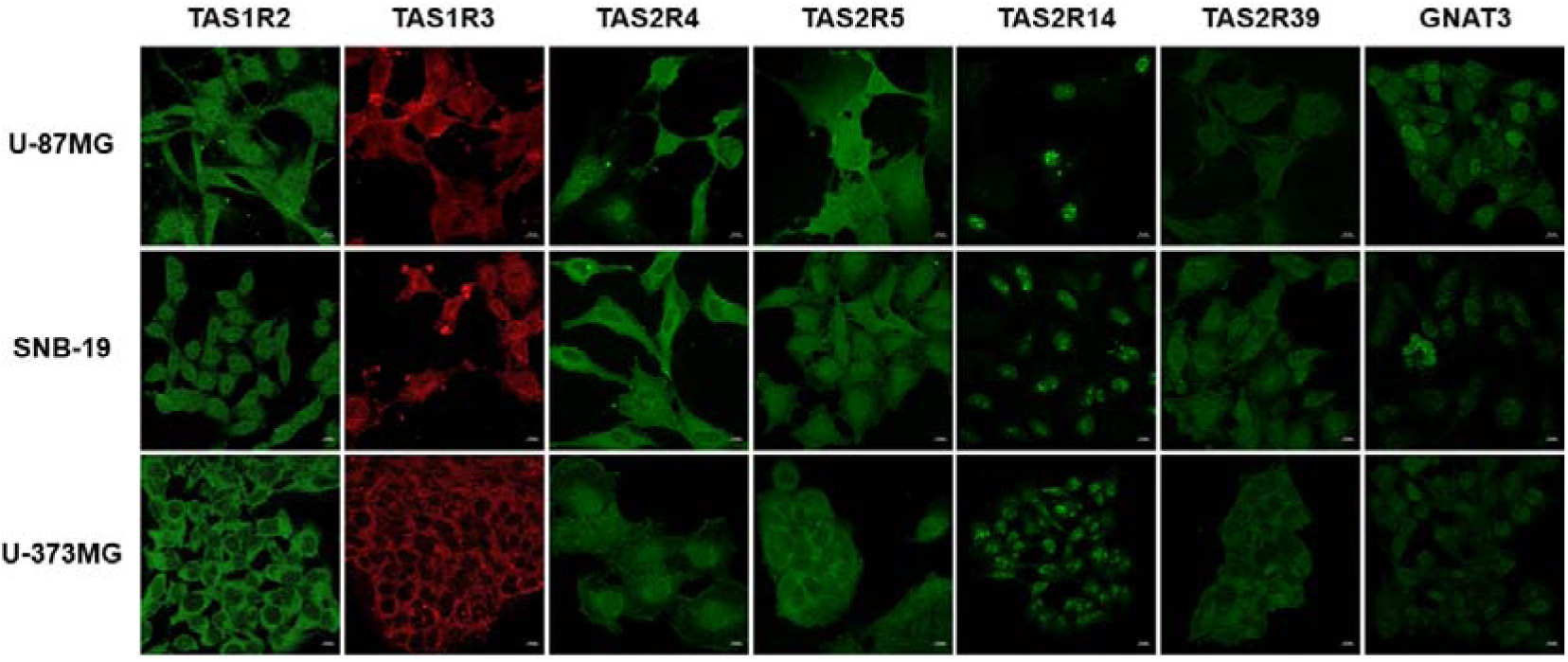
Immunofluorescence detection of sweet and bitter taste receptors, and taste signalling pathway effector GNAT3, in glioblastoma cell lines. Confocal microscopy images of the sweet taste receptor subunits TAS1R2 (green) and TAS1R3 (red), bitter taste receptors TAS2R4, TAS2R5, TAS2R14 and TAS2R39 (green), and taste signalling pathway effector GNAT3 (green) in glioblastoma cell lines (U-87MG, SNB-19, U-373MG). Nuclei were stained with Hoechst 33342. Scale bar: 10 µm.

In addition, all the taste-related proteins were detected at the expected size (Figure 2B). The analysis of the sweet taste receptor subunits TAS1R2 and TAS1R3 relative protein expression is significantly higher in the U-87MG glioblastoma cell line (Figures 2C and 2D). In addition, TAS2Rs relative protein expression revealed that only TAS2R4 expression is significantly higher in human astrocytes than in U-87MG (3.27-fold) or any of the other cell lines (Figure 2E). TAS2R14 expression was about 7-fold higher in U-373MG glioblastoma than in the other cell lines or in human astrocytes (Figure 2G). No statistical differences were observed for TAS2R5 and TAS2R39 protein levels (Figures 2F and 2H).

The subcellular localisation of the sweet taste receptor TAS1R2/TAS1R3, and bitter taste receptors TAS2R4, TAS2R5 and TAS2R39 was mainly cytoplasmatic and in the plasma cell membrane. Interestingly, TAS2R14 was located exclusively at the nucleus of glioblastoma cells (Figure 3). The downstream effector GNAT3 was predominantly found in the cytoplasm of glioblastoma cells but was also detected in the nucleus (Figure 3).

### 3.3 Taste receptors are expressed in human glioblastoma

We analysed the expression of the sweet taste receptor subunits TAS1R2 and TAS1R3, and the four selected TAS2Rs, in paraffin-embedded sections of 10 human glioblastoma samples obtained by surgical resection of the tumour. Taste receptors’ immunoreactivity was observed in all cases (Figure 4), although with receptor-, individual- and region-dependent variations. No immunostaining was seen in sections where primary antibodies were omitted (negative controls). In general terms, taste receptors’ immunoreactivity was found in the cytoplasm of large and medium-sized glioblastoma cells in areas of high cellular density, with weak or no immunoreactivity in less differentiated and cellular density areas. Immunostaining of glioblastoma cells was in striking contrast with the negativity of fibrous regions with predominant fibroblast-like cells and necrosis areas. In addition, TAS2R39 immunoreactivity was lower and more dispersed when compared with the other assessed TAS2Rs (Figure 4).

**Figure 4.**
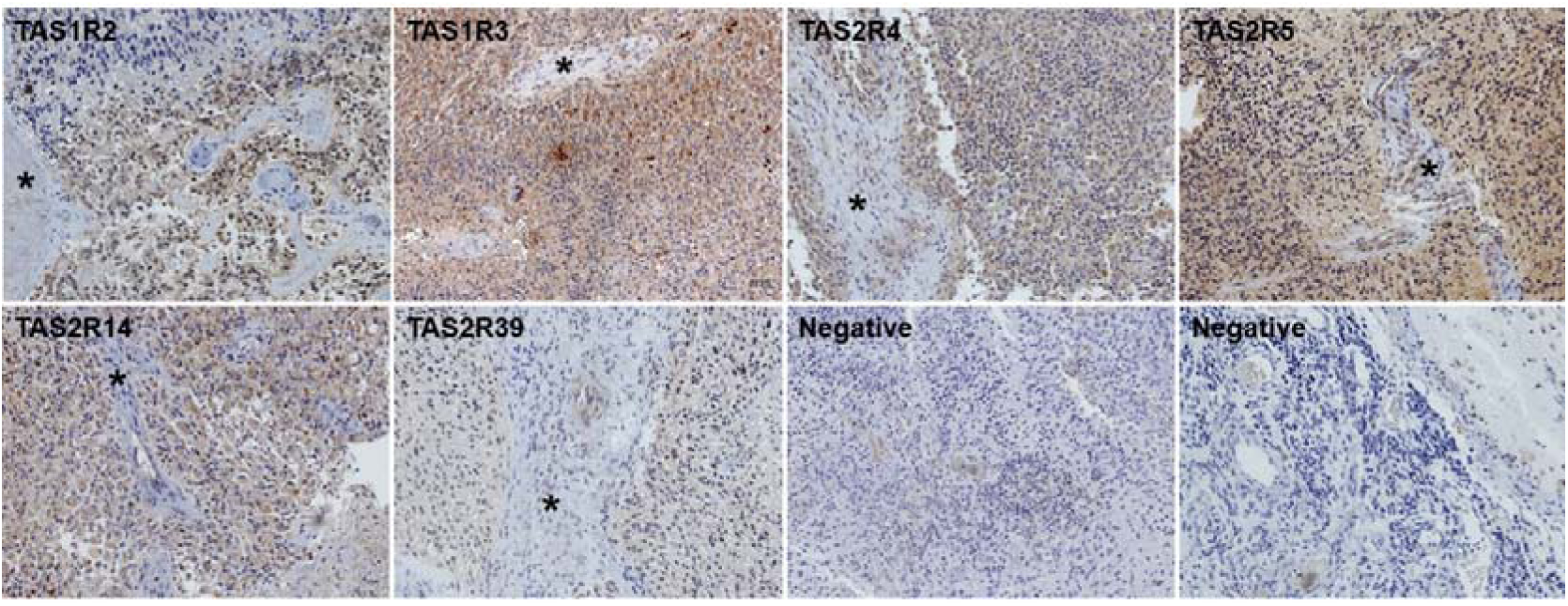
Immunostaining of sweet and bitter taste receptors in tumour samples of human glioblastomas. The sweet taste receptor subunits TAS1R2 and TAS1R3, and bitter taste receptors TAS2R4, TAS2R5, TAS2R14 and TAS2R39 decorate the cytoplasm of large and medium size glioblastoma cells in areas with high cellular density, contrarily to less differentiated or with loose cellularity areas. In the same sections, fibrous areas (asterisks) are negative. No immunostaining was seen in negative control slices where the primary antibodies were omitted. Representative images of paraffin sections, lightly counterstained with Mayer’s haematoxylin. Scale bar: 50 µm.

### 3.4 Denatonium benzoate elicits Ca^2+^ responses in glioblastoma cells

To analyse the functionality of taste receptors, we assessed U-373MG response to denatonium benzoate (DnB; 2.5 and 5 mM) using a Ca^2+^ functional assay (Figure 5) in the presence and absence of probenecid, a known TAS2Rs antagonist. Our results showed that the response of U-373MG glioblastoma cells to 5 mM DnB (ΔF/F0=1.797±0.395) was abolished in the presence of probenecid (ΔF/F0=0.047±0.010).

**Figure 5.**
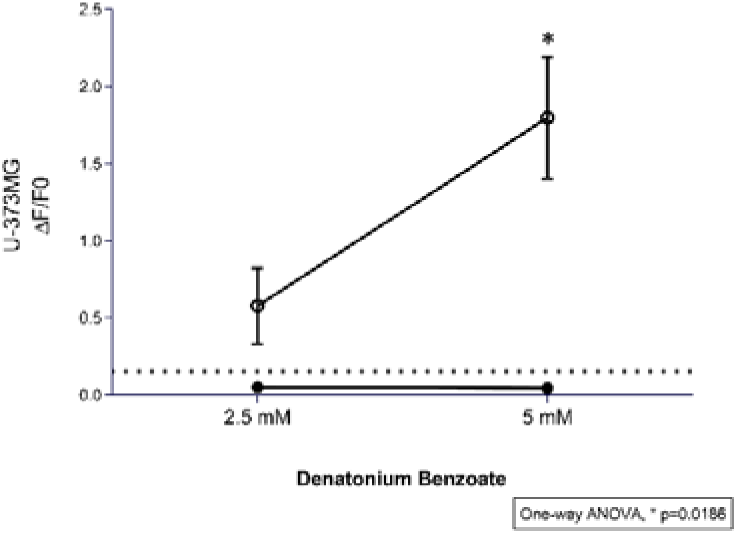
Calcium dose-response curves of U-373MG glioblastoma cells to denatonium benzoate. Dose-response curves of different concentrations of denatonium benzoate (DnB; 2.5 and 5 mM), in the presence (●) or absence (○) of 1 mM probenecid, a known TAS2R inhibitor. Dot line: Ca2+ levels measured in cells with vehicle only (DMSO ≤ 0.5%). Results are presented as mean ± SEM. Statistical analysis was performed by two-way ANOVA followed by Tukey’s multiple comparisons test. [N≥3 independent experiments; *versus probenecid+DnB].

## 4. Discussion

The expression and function of the taste transduction pathway, including both bitter taste receptors (TAS2Rs) and sweet taste receptor (TAS1R2/TAS1R3), have been implicated in several types of cancer, particularly in breast and pancreatic cancer cells, and are associated with chemoresistance and metabolic regulation [27–29]. However, no studies have focused on glioblastoma or any other brain neoplasms despite their high levels of chemoresistance and limited treatment options, as well as reliance on altered metabolic processes such as the Warburg effect, characterised by increased glucose consumption [28,30,31]. Given the potential role of taste receptors in drug response and nutrient sensing, we aimed to profile their expression and functionality in glioblastoma cells.

In this study, we profiled the mRNA expression of 20 of the 26 human TAS2Rs, TAS1R2/TAS1R3 and TAS1R1/TAS1R3 in human astrocytes and three glioblastoma cell lines. The functionality of the taste-signalling pathway was further supported by the detection of the downstream effector protein GNAT3, the G protein specific of the taste-signalling pathway [32], and by Ca^2+^ imaging experiments. Bitter taste receptors TAS2R4, TAS2R5, TAS2R14 and TAS2R39, along with the sweet taste receptor subunits TAS1R2/TAS1R3, were selected to carry out further experiments given their expression in these cell lines, therapeutic relevance, and the availability of validated primary antibodies. Notably, we observed variable levels of both TAS2Rs and TAS1R2/TAS1R3 at the mRNA and protein levels, suggesting post-transcriptional and translational regulation. Previous studies already reported differential levels of TAS2R4 and TAS2R14 in breast cancer compared to normal breast [13,25,33]. Interestingly, TAS2R38 is upregulated in tumour cells from pancreatic cancer patients and tumour-derived cell lines compared to a pancreatic stellate cell line [34]. Other studies reported that mRNA expression of TAS2R14 and TAS2R38 is decreased in both prostate and ovarian cancer cells [35]. In addition, TAS2Rs polymorphic variants, including those of TAS2R4 and TAS2R14, have been studied, but their association with increased cancer risk remains controversial [36,37]. Given the differential expression of TAS2Rs in various cancers, including breast [38,39], pancreatic [34], prostate, and ovarian cancers [35], their role in tumour biology remains complex and potentially depends on the organs undergoing malignant transformation. In addition, we also observed an increased expression of TAS1R2/TAS1R3 in the U-87MG glioblastoma cell line, suggesting that it might be an advantage for tumour development. In cancer cells, including glioblastoma, higher sweet taste receptor expression could enhance glucose sensing and uptake from the tumour microenvironment [40]. This suggests their potential to be used as therapeutic targets in the cells or tissues where they are expressed, underscoring the importance of further studying these receptors in cancer.

Besides identifying the taste receptors expressed in glioblastoma cells, we also analysed their subcellular location. Curiously, TAS2R14 exhibited exclusive nuclear localisation in glioblastoma, whereas cytoplasmic and/or membrane-bound in human astrocytes. This was an intriguing finding whose significance remains unclear, though prior studies have reported nuclear localisation of TAS2R38 in non-ciliated or inflamed airway tissues [41–43], and of TAS2R13 and TAS2R42 in head and neck squamous cell carcinomas [42]. To our knowledge, no systematic studies are addressing the subcellular location of taste receptors in cancer compared to non-cancer cells.

We also investigated the expression patterns of TAS2Rs and TAS1R2/TAS1R3 in human glioblastoma tumour samples. Both bitter and sweet taste receptors were found in high cellular density areas, contrasting with fibrous regions. Similar results were observed by Gaida *et al*. in pancreatic ductal adenocarcinoma tissue samples, where TAS2R38 was stained more prominently in tumour-infiltrating cells and absent in surrounding normal tissue [34]. In addition, TAS2R4, TAS2R5, and TAS2R14 immunoreactivity was higher and less dispersed than TAS2R39, suggesting receptor-specific expression patterns. Given glioblastoma’s high metabolic demands, the TAS1R2/TAS1R3 expression in high cellular density areas may confer an advantage by enhancing glucose sensing and uptake from the tumour microenvironment [40]. In accordance, the sweet taste receptor has been detected in nutrient-sensitive brain regions such as the hypothalamus and choroid plexus, where their expression is influenced by nutritional status [44,45].

Finally, to explore the functionality of taste receptors, particularly TAS2Rs, in glioblastoma cells, we performed a single-cell calcium imaging functional assay with denatonium benzoate (DnB) in the presence or absence of probenecid, a TAS2R antagonist [46,47]. Our findings showed that DnB elicited intracellular Ca^2+^ levels in U-373MG glioblastoma cells in a dose-dependent manner, which were significantly reduced with probenecid, confirming the activation of the bitter taste signalling pathway. Since many TAS2R ligands, including flavonoids, alkaloids, and cannabinoids, have demonstrated anti-proliferative and pro-apoptotic effects in various cancers (reviewed in [48–50]), further research into both TAS2Rs and TAS1R2/TAS1R3 in glioblastoma may reveal novel therapeutic strategies targeting tumour metabolism and drug resistance.

Overall, we provided evidence that the taste-signalling pathway, particularly the bitter and sweet taste receptors, are part of the chemo-sensing apparatus of glioblastoma cells, adding novel players to the crosstalk of these cancer cells with their microenvironment.

## Acknowledgements

This work was supported by the Fundação para a Ciência e Tecnologia (FCT, Portugal) project grants (PTDC/BIM-ONC/7121/2014, UID/Multi/00709/2013 and UID/Multi/00709/2019), by CENTRO 2020 and Lisboa 2020 project grant (POCI-01-0145-FEDER-016822), and FEDER funds through the POCI – COMPETE 2020 – Operational Programme Competitiveness and Internationalization in Axis I – Strengthening research, technological development and innovation (POCI-01-0145-FEDER-007491). Ana R. Costa was recipient of a PhD fellowship (UI/BD/151025/202) funded by FCT through the Portuguese state and EU budgets through the European Social Fund. Ana C. Duarte was recipient of a grant from CENTRO 2020 program through the ICON project (Interdisciplinary Challenges On Neurodegeneration; CENTRO-01-0145-FEDER-000013). We also acknowledge the support of the Portuguese Platform of Bioimaging (PPBI) [PPBI-POCI-01-0145-FEDER-022122] and the resources provided by the Fluorescence Microscopy Unit of RISE-Health, UBI.

