## Supplementary Material for "Taste receptors’ profiling in glioblastoma"

**Table S1.** List of primers used for RT‐PCR mRNA expression for the bitter, sweet and *umami* taste receptors in human astrocytes and glioblastoma cell lines.

| **Ref Seq** | **Gene Name** | **Gene Symbol** | **Primer Sequence 5’ – 3’** | **Size (bp)** | **A.T.**  **(ºC)** |
| --- | --- | --- | --- | --- | --- |
| [NM_138697.4](https://www.ncbi.nlm.nih.gov/gene/80835) | Taste receptor, type 1, member 1 | *TAS1R1* | Fw: GCCTGCAGCTTCTCATTTCC  Rv: AGCTACAAGACCTGTCACACA | 174 | 58 |
| [NM_152232.6](https://www.ncbi.nlm.nih.gov/gene/80834) | Taste receptor, type 1, member 2 | *TAS1R2* | Fw: CTCGGCTGTGACAAAAGCAC  Rv: CCTTGCGGGTCGAAGAAGAT | 119 | 58 |
| [NM_152228.3](https://www.ncbi.nlm.nih.gov/gene/83756) | Taste receptor, type 1, member 3 | *TAS1R3* | Fw: GACAGAGCGCCTGAAGATCC  Rv: CGATGTCGTCTGGGTTTTGC | 173 | 58 |
| [NM_019599.3](https://www.ncbi.nlm.nih.gov/nuccore/NM_019599.3) | Taste receptor, type 2, member 1 | *TAS2R1* | Fw: TCCCAGCTGTCTGAAGGTGT  Rv: GCCTGAAGGGGACATGTTGTA | 1334 | 60 |
| [NM_016943.2](https://www.ncbi.nlm.nih.gov/nuccore/NM_016943.2) | Taste receptor, type 2, member 3 | *TAS2R3* | Fw: ATCAGGGCTGCCTAATTGCT  Rv: GTCCTGTAGTCTTGAGCCAGG | 1035 | 60 |
| [NM_016944.2](https://www.ncbi.nlm.nih.gov/nuccore/NM_016944.2) | Taste receptor, type 2, member 4 | *TAS2R4* | Fw: TGTTTCGGTTATTCTATTTCTCTGC  Rv: CCTGGAGAGTAAAGGGTGGC | 823 | 58 |
| [NM_018980.3](https://www.ncbi.nlm.nih.gov/nuccore/NM_018980.3) | Taste receptor, type 2, member 5 | *TAS2R5* | Fw: ACTACCAGGGGATCTGACCTC  Rv: CCGAGCACACACTGTCTTCC | 937 | 60 |
| [NM_023919.2](https://www.ncbi.nlm.nih.gov/nuccore/NM_023919.2) | Taste receptor, type 2, member 7 | *TAS2R7* | Fw: GCAGGTGTGGATGTCAAACTC  Rv: ATGACTTGAGGGGTAGATTAGAGC | 900 | 60 |
| [NM_023918.3](https://www.ncbi.nlm.nih.gov/nuccore/NM_023918.3) | Taste receptor, type 2, member 8 | *TAS2R8* | Fw: TGTTCAGTCCTGCAGATAACATC  Rv: GCATTCTGACAAATGTCTGCC | 897 | 58 |
| [NM_023917.2](https://www.ncbi.nlm.nih.gov/nuccore/NM_023917.2) | Taste receptor, type 2, member 9 | *TAS2R9* | Fw: GGCATGCCAAGTGCAATAGAG  Rv: AGGGGTCTCTATGGAACAAAAGG | 950 | 58 |
| [NM_023921.1](https://www.ncbi.nlm.nih.gov/nuccore/NM_023921.1) | Taste receptor, type 2, member 10 | *TAS2R10* | Fw: GCTACGTGTAGTGGAAGGCA  Rv: TGCAGTACCCTCAAAGAGGC | 876 | 60 |
| [NM_023920.2](https://www.ncbi.nlm.nih.gov/nuccore/NM_023920.2) | Taste receptor, type 2, member 13 | *TAS2R13* | Fw: GCTAGGGCTCAGCAGAGAAAT  Rv: GGCAAGTCCAAACTTCCCTAAT | 1607 | 58 |
| [NM_023922.1](https://www.ncbi.nlm.nih.gov/nuccore/NM_023922.1) | Taste receptor, type 2, member 14 | *TAS2R14* | Fw: TGGGTGGTGTCATAAAGAGCAT  Rv: CTGAGGGCTCCCCATCTTTG | 924 | 58 |
| [NM_016945.3](https://www.ncbi.nlm.nih.gov/nuccore/NM_016945.3) | Taste receptor, type 2, member 16 | *TAS2R16* | Fw: GTCCAGGAAGACACTTTGGAGT  Rv: TAGGCCTAGCACTTTCCCTT | 909 | 60 |
| [NM_176817.5](https://www.ncbi.nlm.nih.gov/nuccore/NM_176817.5) | Taste receptor, type 2, member 38 | *TAS2R38* | Fw: TTTCTGCACTGGGTGGCAA  Rv: GGCATATTTATGAAGACTCACAGGC | 1141 | 60 |
| [NM_176881.2](https://www.ncbi.nlm.nih.gov/nuccore/NM_176881.2) | Taste receptor, type 2, member 39 | *TAS2R39* | Fw: TCTGCGATCCTGCAGAAAGT  Rv: GATGAAGTCGAAGCTGAAGCC | 930 | 58 |
| [NM_176882.2](https://www.ncbi.nlm.nih.gov/nuccore/NM_176882.2) | Taste receptor, type 2, member 40 | *TAS2R40* | Fw: TCTTGGCGCAGAAACCTGAA  Rv: TTCCAGTCACAGAGTCTGCC | 1015 | 58 |
| [NM_176883.2](https://www.ncbi.nlm.nih.gov/nuccore/NM_176883.2) | Taste receptor, type 2, member 41 | *TAS2R41* | Fw: GCAGCGAATGGCTTCATTGT  Rv: AACAGGAGCTGCGAGAACAC | 833 | 60 |
| [NM_181429.2](https://www.ncbi.nlm.nih.gov/nuccore/NM_181429.2) | Taste receptor, type 2, member 42 | *TAS2R42* | Fw: ATGGCCACCGAATTGGACA  Rv: CTACAAAGGTAAAGGGTTTGGTGT | 945 | 58 |
| [NM_176884.2](https://www.ncbi.nlm.nih.gov/gene/259289) | Taste receptor, type 2, member 43 | *TAS2R43* | Fw: TATCTGGGCAGTGATCAACCA  Rv: CCCCAACAACATCACCAGAATG | 148 | 56 |
| [NM_176885.2](https://www.ncbi.nlm.nih.gov/nuccore/NM_176885.2) | Taste receptor, type 2, member 44 | *TAS2R44* | Fw: TTTTTCCAGTGTGGTAGTGGTTCT  Rv: GATGAAGGCTTCTCTCCTTTCACC | 900 | 58 |
| [NM_176888.2](https://www.ncbi.nlm.nih.gov/nuccore/NM_176888.2) | Taste receptor, type 2, member 48 | *TAS2R48* | Fw: GAACAAGTGTTACTAAGCCTGC  Rv: CTTCTTTCACTCAGCGTGTCA | 952 | 58 |
| [NM_176890.2](https://www.ncbi.nlm.nih.gov/nuccore/NM_176890.2) | Taste receptor, type 2, member 50 | *TAS2R50* | Fw: ACAACCAGTGATATTAGGCTTGC  Rv: TCAGGTCTTTTACTCAGCACCT | 963 | 58 |
| [NM_177437.1](https://www.ncbi.nlm.nih.gov/nuccore/NM_177437.1) | Taste receptor, type 2, member 60 | *TAS2R60* | Fw: TCCTTTTACGCCTGGTAGCAA  Rv: AGGAACGACGACTCTTCAGC | 864 | 60 |
| Fw – Forward; Rv – Reverse; A.T. – Annealing temperature. | | | | | |

**Table S2.** List of primers used for real‐time quantitative RT-PCR for the bitter and sweet taste receptors in human astrocytes and glioblastoma cell lines.

| **Ref Seq** | **Gene Name** | **Gene Symbol** | **Primer Sequence 5’ – 3’** | **Size (bp)** | **A.T.**  **(ºC)** |
| --- | --- | --- | --- | --- | --- |
| [NM_016944.2](https://www.ncbi.nlm.nih.gov/nuccore/NM_016944.2) | Taste receptor, type 2, member 4 | *TAS2R4* | Fw: CTGGAATCCCCAGACGGAAG  Rv: CTGGACCAGGGTAGCAACTG | 97 | 60 |
| [NM_018980.3](https://www.ncbi.nlm.nih.gov/nuccore/NM_018980.3) | Taste receptor, type 2, member 5 | *TAS2R5* | Fw: CTTTTCCAGAGCAGCCGTTG  Rv: GTGGCAAACCATAAGCTGGC | 80 | 60 |
| [NM_023922.1](https://www.ncbi.nlm.nih.gov/nuccore/NM_023922.1) | Taste receptor, type 2, member 14 | *TAS2R14* | Fw: GGTGAACTGTATTGACTGGGTC  Rv: GCTGGGAAAAACACAGACACAC | 135 | 58 |
| [NM_176881.2](https://www.ncbi.nlm.nih.gov/nuccore/NM_176881.2) | Taste receptor, type 2, member 39 | *TAS2R39* | Fw: ATGTGGTCGGTCTGGCTTTT  Rv: TGCTTCCCATGTGTAGGGTG | 120 | 58 |
| [NM_001357943.2](https://www.ncbi.nlm.nih.gov/nucleotide/1676440496) | Glyceraldehyde 3-phosphate dehydrogenase | *GAPDH* | Fw: ATGGGGAAGGTGAAGGTCG  Rv: GGGGTCATTGATGGCAACAATA | 108 | 58 / 60 |
| Fw – Forward; Rv – Reverse; A.T. – Annealing temperature. | | | | | |

**Table S3.** List of antibodies used for Western blot, immunohistochemistry, and immunofluorescence of bitter and sweet taste receptors’ protein detection.

|  | **Host** | **Catalog Number** | **WB** | **IHC** | **IF** |
| --- | --- | --- | --- | --- | --- |
| *Primary Antibodies* | | | | | |
| TAS1R2 | Rabbit | SantaCruz Biotechnology #sc-50306 | 1:250 | – | 1:100 |
| TAS1R2 | Rabbit | ThermoFisher Scientific #PA5-34254 | – | 1:500 | – |
| TAS1R3 | Goat | SantaCruz Biotechnology #sc-22458 | – | – | 1:100 |
| TAS1R3 | Rabbit | ThermoFisher Scientific #PA5-102260 | 1:1000 | 1:2000 | – |
| TAS2R4 | Rabbit | ThermoFisher Scientific #OSR00153W | 1:500 | 1:1500 | 1:500 |
| TAS2R5 | Rabbit | ThermoFisher Scientific #OSR00154W | 1:500 | 1:1500 | 1:500 |
| TAS2R14 | Rabbit | ThermoFisher Scientific #PA5-39710 | 1:300 | 1:750 | 1:300 |
| TAS2R39 | Rabbit | ThermoFisher Scientific #PA5-39711 | 1:300 | 1:500 | 1:300 |
| TAS2R43 | Rabbit | ThermoFisher Scientific #PA5-103257 | – | – | 1:250 |
| β-Actin | Mouse | Sigma-Aldrich #A1978 | 1:30000 | – | – |
| GNAT3 | Rabbit | SantaCruz Biotechnology #sc-395 | 1:250 | – | 1:100 |
| GNAT3 Peptide | Rabbit | SantaCruz Biotechnology #sc-395 P | 1:250 | – | – |
| *Secondary Antibodies* | | | | | |
| Alexa Fluor^®^ 488 Anti-Rabbit | Goat | ThermoFisher Scientific #A-11008 | – | – | 1:1000 |
| Alexa Fluor^®^ 647 Anti-Goat | Chicken | ThermoFisher Scientific #A-21469 | – | – | 1:1000 |
| Anti-rabbit IgG-HRP | Goat | ThermoFisher Scientific #31466 | 1:20000 | – | – |
| Anti-mouse IgG-HRP | Goat | SantaCruz Biotechnology #sc-2005 | 1:30000 | – | – |
| IF – Immunofluorescence; IHC – Immunohistochemistry; WB – Western blot. | | | | | |
